# Gill-associated microbiome as indicators of population stress and condition in Eastern Baltic Cod

**DOI:** 10.64898/2025.12.16.694594

**Authors:** Maria Cortazar Chinarro, Maddi Garate-Olaizola, Johanna L Fröjd, Yvette Heimbrand, Jane W. Behrens, Anssi Laurila

## Abstract

The skin microbiome contributes to host health by facilitating pathogen exclusion, priming the immune system modulation and maintaining skin homeostasis. We examined how the composition of the skin microbiota is associated with body size and condition in two populations of eastern Baltic cod (*Gadus morhua callarias*), at Bornholm basin (ICES subdivision 25) and in the Åland Sea (subdivision 29). These locations are characterized by differences in body size and experience distinct hydrographic conditions, including salinity, oxygen and temperature. We used 16s rRNA gene metabarcoding to compare skin microbiota diversity and composition in these populations. We found significant differences between the populations in both alpha and beta diversity, with higher microbial diversity observed in the Åland cod characterized by larger body size. Moreover, we identified a greater number of significantly enriched Gene Ontology (GO) metabolic pathways in Åland cod as compared to those from the Bornholm Basin, with many pathways absent in Bornholm. However, other GOs, such as those associated with the nervous system, were uniquely detected in the Bornholm cod. Differences in microbial communities can be influenced by external environmental factors, determined by the habitats where individual cod reside. These results indicate a potential relationship between the skin microbiome and host health; however, direct evidence for a causal impact remains limited, highlighting the importance of incorporating host skin microbiota interactions in studies of aquatic populations.

## Introduction

A key question in ocean microbial ecology is what drives contemporary biogeographic patterns of microbial diversity in the benthic and pelagic as well as host-associated microbial communities (Delleuze et al., 2024, Langenheder & Lindström, 2019). Historical biogeographic patterns of microbial diversity are influenced by the legacy of past events and conditions—such as disturbances, colonization sequences, and environmental fluctuations—and contemporary environmental selection pressures (drift selection, speciation and dispersal), playing a pivotal role in shaping microbial community composition in the oceans (Martiny et al., 2006, Fuhrman, 2009). These processes also influence the assembly of host-associated microbial communities, affecting their diversity and biogeographical patterns (Kohl, 2020, Vellend, 2010).

Numerous studies have characterized host-associated microbiomes and explored population-level variation in gut microbial communities (Truong et al., 2017, Couch & Epps, 2022). Generally, host-associated interface microbiomes in direct contact with external environments, such as, skin or gills, have been largely overlooked. Only a few studies have examined their biogeographical distribution and the ecological factors that shape these patterns. For example, (Morelan et al., 2019) demonstrated that microbial communities associated with temperate sea anemones (*Anthopleura elegantissima*) exhibited only weak links to biogeographic patterns. In contrast, microbial communities in the Californian mussel (*Mytilus californianus*) showed a strong compositional differentiation along a latitudinal gradient (Neu et al., 2021). Research on the biogeographical patterns of host-associated microbiomes across populations remains limited, representing a critical gap in our understanding of microbial diversity and its ecological drivers in aquatic systems.

Aquatic environments harbor distinct and often more diverse bacterial communities compared to terrestrial environments. This diversity is shaped by environmental conditions such as pH, salinity, temperature, and nutrient availability (Wang et al., 2023). In fish, the key site for host–microbe interactions is the skin mucus layer, which serves as the primary barrier and interfaces with the aquatic environment. The skin mucus microbiota is therefore highly responsive to environmental changes (Krotman et al., 2020). Its composition is shaped not only by surrounding conditions, but also by host-associated factors such as genetic background (Chiarello et al., 2015) and anthropogenic disturbances, including eutrophication (Côte et al., 2022) and pollution (Mlejnková & Sovová, 2010). Here, fish skin mucus acts as an important protective barrier against external pathogens and parasites, composed of glycoproteins (mucins), innate immune cells, lipids, and a diverse microbial community (Esteban & Cerezuela, 2015). The associated microbial assemblage is typically dominated by Proteobacteria, its composition differing markedly from that of the surrounding water column (Gomez & Primm, 2021).

The Baltic cod (*Gadus morhua callarias*) is an ecologically and evolutionarily diverged subspecies of the Atlantic cod. It is divided in two stocks of which the eastern stock, eastern Baltic cod, has suffered from high fishing pressure, high infection load with the parasitic liver worm *Contracaecum osculatum* and deteriorating oxygen conditions, the last factor resulting in spawning habitat deterioration, reduction in benthic prey availability and direct physiological effects (Plambech et al., 2013, Sokolova et al., 2018, Ryberg et al., 2020, Eero et al., 2015, Eero et al., 2023). Together these have led to poor nutritional status and reduced growth which led to complete closure of the fishery in 2019 (Mion et al., 2021, Bryhn et al., 2022) ICES 2025). Currently, only one of the three historical spawning areas of eastern Baltic cod —the Bornholm Basin—remains an active site of reproduction as two of these areas have become hypoxic due to the absence of saltwater inflows from the Atlantic, leading to declining salinity and oxygen concentrations (Hinrichsen et al., 2016, Orio et al., 2019). A recent study investigating growth trends in eastern Baltic cod over 25 years of intensive fishing reported a 48% decline in body size and reduced growth rate, particularly among individuals from southern locations, between 1996 and 2019 (Han et al., 2025, ICES, 2021). However, the cod in the Åland Sea at the northern Baltic proper are much larger than the cod in the southern Baltic Sea, yet it remains unclear if this is because of differences in age structure or if the fish in the Åland Sea have superior growth rate (Heimbrand et al., 2023, Bergström et al., 2025). Finally, high infection loads of *C. osculatum* is associated with decreased physiological condition of the cod, depressed energy turnover and reduced the digestive organ masses (Ryberg, 2020), all related to infection-induced severe inflammation in the liver tissue and reduction of hepatocyte fat content (Behrens et al., 2023). These pathological effects seem irreversible even when access to food is plentiful, and may directly impact cod also by impairing swimming performance and foraging efficiency, ultimately constraining growth in eastern Baltic cod (Behrens et al., 2025).

In this study, we investigated gill-associated microbiomes in eastern Baltic cod from two areas in the Baltic Sea, the Bornholm Basin (ICES subdivision 25) and the Åland Sea (subdivision 29). These areas differ in several environmental aspects, including salinity, temperature, and oxygen levels. For example, cod in the Åland Sea inhabit waters of 5–7.5 psu salinity, while those in the Bornholm Basin experience higher salinity (7–17 psu) (Westerlund et al., 2021, Snoeijs-Leijonmalm & Andrén, 2017). We analyzed the 16S rRNA bacterial profiles, predicted GO metabolic pathways and investigated whether differences in body size and condition between the two areas were associated with gill-associated microbiota composition, diversity and liver worm infectiond. We expect a reduction in bacterial diversity in the Åland population compared to the Bornholm population, aligning with previous studies that have documented a decline in microbial diversity with increasing latitude in aquatic ecosystems (Milici et al., 2016, Barton et al., 2010). Furthermore, bacterial composition is expected to differ markedly between habitats, reflecting e.g. variation in oxygen, temperature and salinity conditions. As strong parasite infection can be associated with microbiome dysbiosis, we expected altered or low microbial diversity in individuals heavily infected by *C. osculatum* liver worms.

## Methods

### Sample collection

Eastern Baltic cod were caught with bottom trawl in ICES Subdivision 25 originated in the Bornholm basin (n = 32) and with gillnets in the northern population of Subdivision 29 (Åland sea, n = 27) in November 2023. We swabbed the gill filaments using rayon swabs (MWN: 100mwn). We determined sex of each individual and measured total length and standard deviation from the mean (TL: Bornholm; 318 ± 47 mm, Åland; 691 ± 162 mm), standard weight (SW: Bornholm; 291 ± 143 g, Åland; 3268 ± 2666 g), liver weight (LW: Bornholm; 12.39 ± 7.36 g, Åland; 178.51 ± 130 g), and counted the number of nematodes on the surface of liver. We assumed that all worms on the surface of liver nematodes were *C. osculatum* as previous studies have shown that close to 100 % of cod liver nematodes in the Baltic Sea belong to this species (Ryberg, 2020, Sokolova, 2013).

### DNA extraction and metabarcoding 16s library preparation

Bacterial community DNA was extracted from gill swabs using the DNeasy PowerSoil Kit (Qiagen), following the manufacturer’s protocol. The extracted DNA was quantified using both a spectrophotometer (NanoDrop; Thermo Fisher Scientific) and a fluorometric assay (Qubit; Invitrogen). Bacterial DNA from the gill area was subjected to 16S rRNA gene amplicon sequencing using the Illumina MiSeq platform (Illumina Inc.). Library preparation followed a two-step PCR protocol. In the first PCR (30 cycles), the hypervariable V4 region of the bacterial 16S rRNA gene was amplified using primers 515F (5′-GTGCCAGCMGCCGCGGTAA-3′) and 806R (5′-GGACTACHVGGGTWTCTAAT-3′) as described in (Cortazar-Chinarro et al., 2024). The primers were modified at the 5′-ends with Illumina adapter sequences to enable downstream indexing. In the second PCR (20 cycles), unique dual indices integrated into the Phusion primers were added to both ends of the amplicons to individually barcode each sample (see (Cortazar-Chinarro et al., 2024)). The barcoding primers included Illumina sequencing handles to facilitate attachment of the amplicons to the flow cell during sequencing. Both PCR reactions were performed using Phusion High-Fidelity DNA Polymerase (Thermo Fisher Scientific), with PCR mixtures prepared according to the manufacturer’s instructions and supplemented with 20 mg/mL BSA (Bovine Serum Albumin; Thermo Fisher Scientific). In case of contamination, all PCRs (PCR1 and PCR2) were repeated until negative controls showed no amplification.16s Amplicons were purified after each PCR step using the Agencourt AMPure XP purification kit (Beckman Coulter Inc.). Fragment size and DNA concentration were assessed using a Bioanalyzer (Agilent Technologies) and a fluorescence microplate reader (Ultra 384; Tecan Group Ltd.), with quantification performed using the Quant-iT PicoGreen dsDNA Assay Kit (Invitrogen). Equimolar quantities of the purified amplicons were pooled, and sequencing was performed on the Illumina MiSeq platform at NGI/SciLifeLab, Uppsala, Sweden.

### 16s bioinformatic workflow

Raw sequence data were processed using the DADA2 pipeline (Callahan et al., 2016). Forward and reverse reads were trimmed to 240 bp and 200 bp, respectively, using default quality filtering parameters. Amplicon error correction, chimera removal, and paired-end read merging were also performed using DADA2 defaults. Amplicon Sequence Variants (ASVs) were inferred and taxonomically assigned using the SILVA 16S rRNA gene reference database (version 2019; (Henderson et al., 2019). We removed ASVs that not could be taxonomically classified according to (Costa et al., 2022, Couch et al., 2021). Singleton ASVs were filtered out to minimize spurious results. An additional filtering step was applied to remove all ASVs assigned to mitochondria, chloroplasts, or unclassified/uncultured bacteria. This was performed using the *subset_samples*() and *prune_taxa*() functions from the phyloseq R package (McMurdie & Holmes, 2019). Functional profiles were predicted from 16S rRNA gene data using PICRUSt2 (Douglas et al., 2020), and KEGG pathway annotations were obtained via the KEGGREST package in R (Tenenbaum et al., 2019).

### Analysis of bacteriome relative abundance and diversity

To account for uneven sequencing depth across samples, the data were rarefied prior to estimating relative abundance, alpha diversity, and beta diversity metrics. Rarefaction was performed using the *rarefy_even_depth()* function from the phyloseq R package (McMurdie & Holmes, 2019), with a standardized depth of 69,722 reads per sample, determined based on sample distribution and retention rate. For relative abundance analyses, taxa were aggregated at the phylum level, and abundance values were normalized using the *transform_sample_counts*() function in phyloseq (McMurdie & Holmes, 2019). Alpha diversity was assessed using Shannon, Observed Richness, Simpson, and Chao1 indices. Statistical comparisons between populations (Åland vs. Bornholm) were conducted using either Wilcoxon rank-sum or Kruskal–Wallis test, according to their non-parametric data distribution. These analyses were implemented using the *vegan* and *ape* packages in R (Oksanen et al., 2013, Paradis et al., 2019).

We explored phylogenetic relationships among 16S microbial taxa and their distribution in the host populations using hierarchical clustering, performed with the *hclust* function and Ward’s method in R (Kolde & Kolde, 2015). Differences in bacterial community composition between the populations and sexes were evaluated using three complementary approaches. First, we applied PERMANOVA (Permutational Multivariate Analysis of Variance) to assess whether overall microbial community composition significantly differed between groups. Second, to verify the assumption of homogeneity of group dispersions, we performed a PERMDISP (Permutational Analysis of Multivariate Dispersions) test. Third, we conducted an ANOSIM (Analysis of Similarities) to evaluate differences in community composition based on ranked dissimilarities between sampling locations, providing a complementary perspective to the PERMANOVA results. All tests were conducted using multiple distance metrics, including Bray–Curtis, unweighted UniFrac, weighted UniFrac, and Jaccard dissimilarities using the *vegan* package in R.

### Microbial diversity, parasite infection and host traits

We fitted generalized linear models (GLM) to investigate the relationship between alpha diversity (Shannon diversity index) and host phenotypic traits (weight, length, liver weight, number of liver worms sex and condition) using the car package in R (Fox et al., 2012). Liver weight was standardized by dividing it by individual total length to account for body size, resulting in the variable sizeRatio_LW_L. Prior to model fitting, we assessed multicollinearity among predictors to exclude highly correlated variables from the final model. The Shannon diversity index was used as the response variable, with weight, length, sizeRatio_LW_L, and sex included as explanatory variables.To further explore potential trait–diversity relationships independent of population effects, we conducted separate correlation analyses (spearman rank correlations) between the Shannon index and each predictor variable within each population . Additionally, we tested whether hepatosomatic (Nikol′skiĭ, 1963) and Fulton condition (Ricker, 1975) were correlated with shannon bacterial diversity. We also conducted redundancy analyses to examine whether host phenotypic traits were associated with bacterial composition across sampling locations. To assess whether sampling location affected our results, we first quantified variance using a simplified model with population as the fixed predictor and ASVs as the response variable. We then performed variation partitioning using the *varpart()* function in the vegan package to determine the proportion of variance attributable to population versus other variables (Oksanen et al., 2013). We then conducted separate RDA analyses by using the vegan package R for each population to evaluate potential independent associations within each sampling location (Oksanen et al., 2013). Also, we tested for interaction between population and host factors and run partial RDA to control for site effects.

### Differential abundance of taxa and predicted metabolic pathways (KEGG orthologs)

To investigate whether specific bacterial taxa differed in abundance between populations and sexes, we applied differential abundance analyses using DESeq2 with Phyloseq R package and the LindA method implemented in the MicrobiomeSTAT R package (Zhang et al., 2023, McMurdie & Holmes, 2019). For LindA method, non-rarefied count data were transformed into compositional data using the centered log-ratio (CLR) transformation, following the default settings. Additionally, we investigated differences in predicted functional metabolic pathways (KEGG pathways) between populations and sexes using two independent approaches: DESeq2 and LindA, as implemented in the ggpicrust2 package in R (Yang et al., 2023). The KO metabolic pathways panel was generated by Picrust2 softwares (Douglas et al., 2020). Population and sex were used as predictor variables in two separate models. To control for false discovery rate, we applied the Benjamini-Hochberg method for p-value adjustment.

## Results

### 16s bacteriome characterization and diversity

A total of 9,843,238 sequencing reads were obtained from 59 eastern Baltic cod samples for characterizing of the gill-associated bacteriome. Prior to data filtering, 21,285 taxa were identified across nine taxonomic ranks. Following the removal of chloroplast, mitochondrial, and unclassified taxa at any rank, 9677 bacterial taxa remained for downstream analyses. The most abundant phyla were Proteobacteria (45.8% of the total number of sequences), Bacteroidetes (16%), Actinobacteria (12.1%), Firmicutes (11.3 %), Acidobacteria (2.7%) and Verrucomicrobiota (2 %). The rest of the phyla represents less than 5% of the total number of reads: Planctomycetes (1.9 %), Chloroflexi (1.7%), Spirocahetes (0.7%), Fusobacterai (0.7%). Proteobacteria was the most abundant phylum in both populations, exhibiting a slightly higher relative abundance in Bornholm (48.3%, N = 239 phyla) compared to Åland (45.1%, N = 351 phyla; Figure 1; Figure S1).

**Figure 1.**
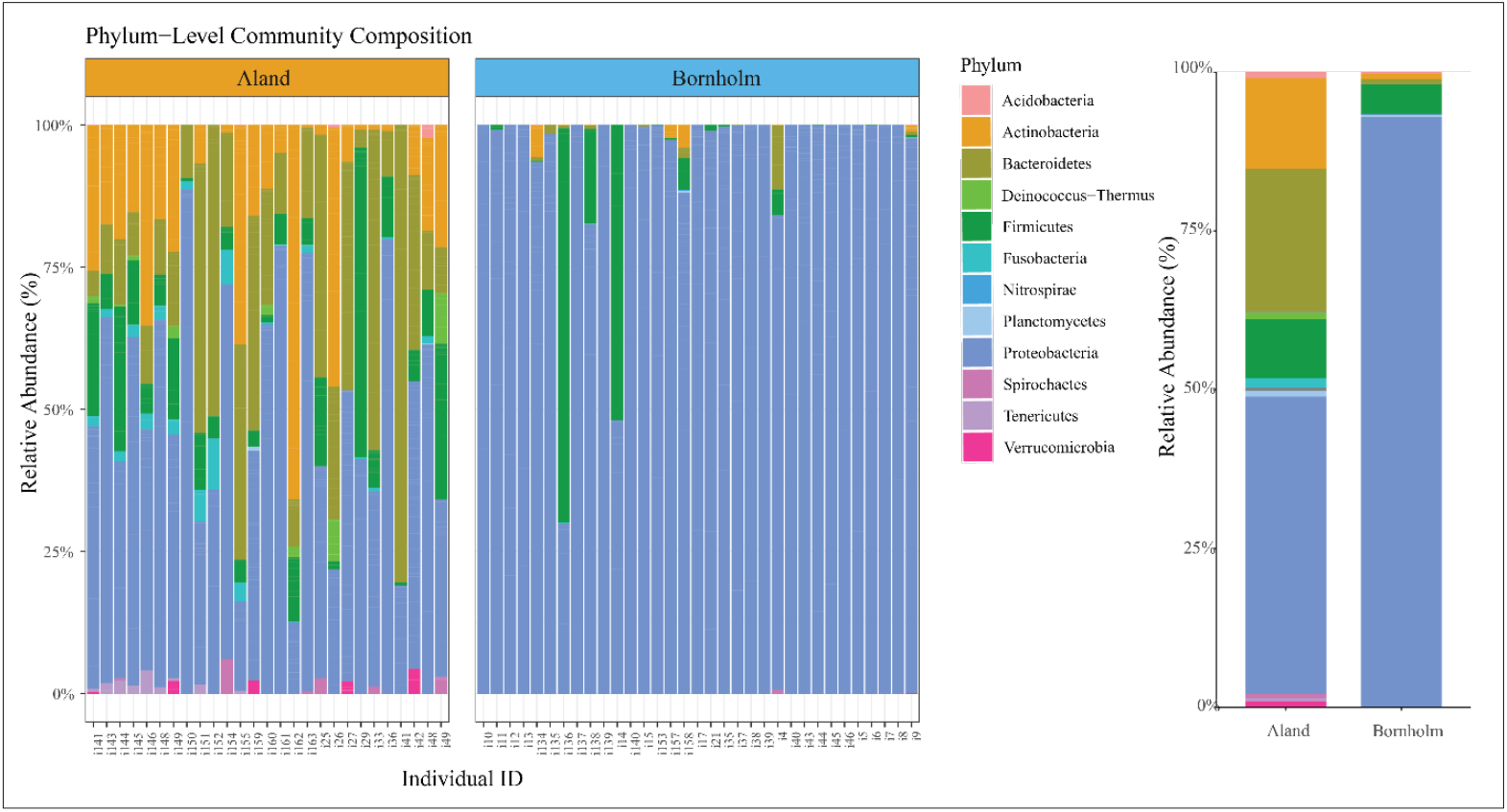
Relative abundance plot comparing individual and population-level bacterial phyla in the gill-associated microbiome between Åland (SD 29) and Bornholm (SD25) locations, highlighting the most prevalent bacterial Phylum.

We detected significantly higher alpha diversity in cod from the Åland Sea than in Bornholm in all diversity metrics (Shannon, Simpson, Chao1 and Observed; p<0.0001; Figure 2). No significant difference in diversity was found between sexes (Åland; DF=1, F=0.064, p=0.80, Bornholm; DF=1, F=0.07, p=0.89). Bacterial community composition differed significantly between Åland and Bornholm in all three statistical approaches: PERMANOVA (R^2^ = 0.143, F = 9.68, p < 0.0001), PERMDISP (DF = 1, F = 12.89, p < 0.001), and ANOSIM (R^2^ = 0.369, p = 0.001). This pattern was further supported by multivariate statistical analysis using Principal Coordinates Analysis (PCoA) based on Bray-Curtis dissimilarity (Figure 3). Hierarchical clustering of 16S bacterial composition revealed distinct population-specific groupings of gill-associated microbiota in cod from Åland and Bornholm, while also indicating potential microbial interchange or connectivity between populations (Figure 3).

**Figure 2.**
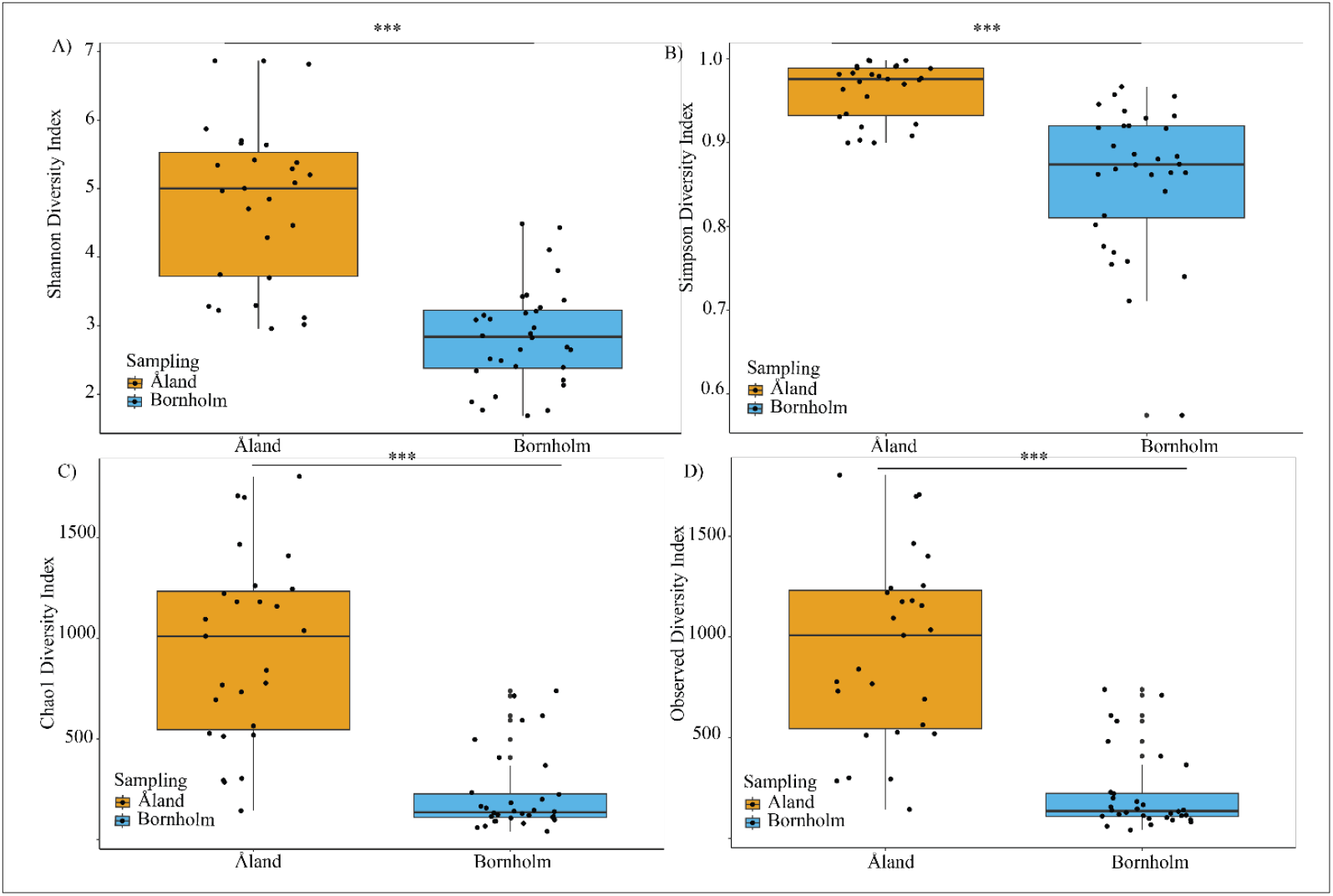
Comparison of alpha diversity indices in the gill microbiomes of Eastern Baltic cod from Åland (orange) and Bornholm (blue). (A) Shannon diversity index, (B) Simpson diversity index, (C) Chao1 diversity index, and (D) observed diversity index, each showing significantly greater diversity in Åland compared to Bornholm (p < 0.001).Boxplots display the range, median, and individual sample diversity values for each site.Alpha diversity comparisons of cod gill microbiomes between Åland (orange) and Bornholm (blue) sampling sites.(A) Shannon index, (B) Simpson index, (C) Chao1 index, and (D) Observed diversity index, with all metrics significantly higher in Åland than Bornholm (p < 0.001). Significant p-values are represented with a (*)

**Figure 3.**
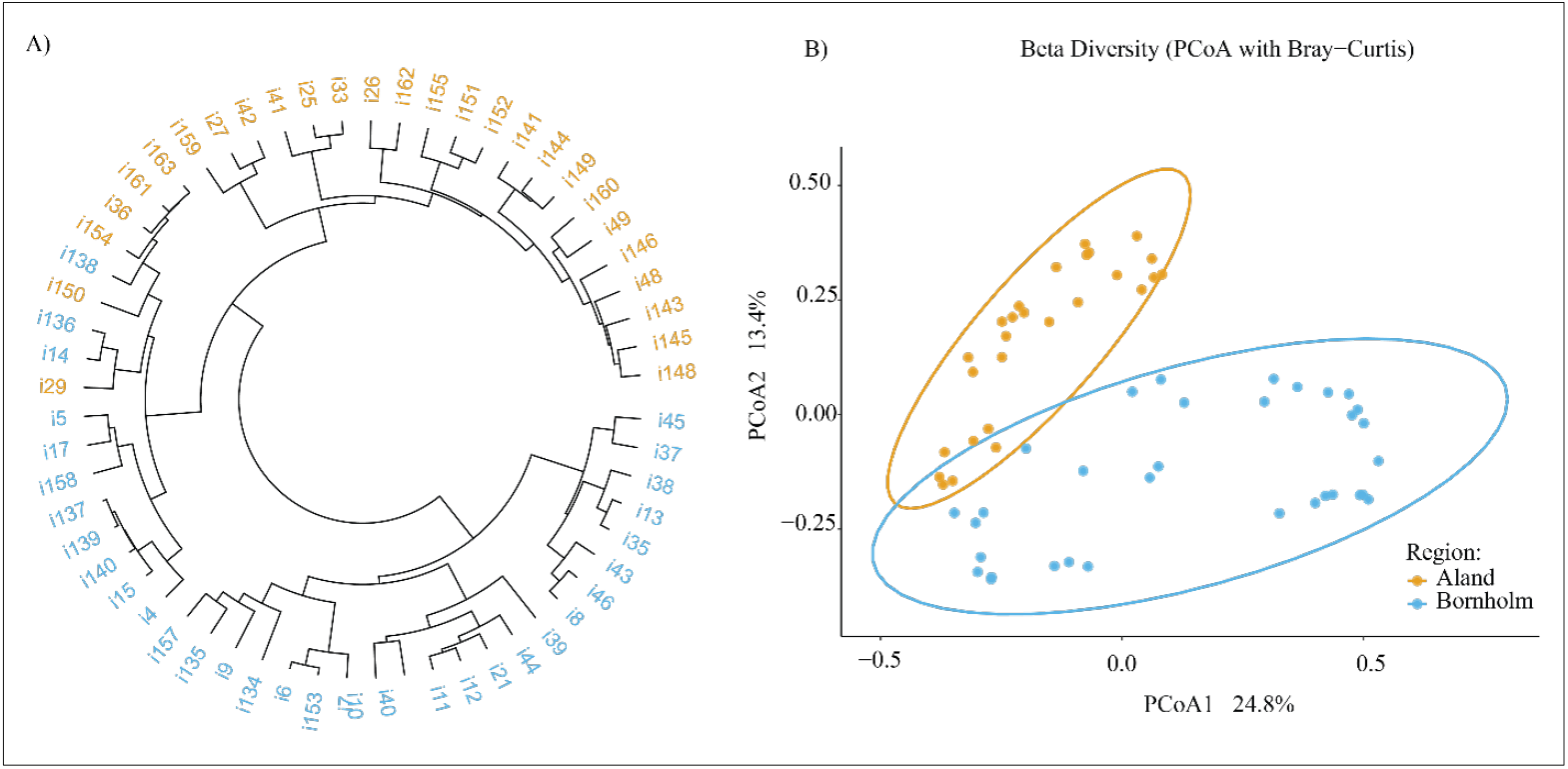
Microbial community structure of Eastern Baltic cod gill samples from Bornholm (blue) and Åland (orange). (A) Hierarchical clustering dendrogram reflecting relationships among individual samples, colored by population. (B) Principal Coordinates Analysis (PCoA) of beta diversity (Bray– Curtis), showing separation between Bornholm and Åland samples at both individual and population levels.

### Microbiome and host traits

Our data reveal larger mean size and higher parasite loads in Åland cod compared to Bornholm cod (Figure S2). Populations showed no significant differences in Fulton condition or hepatosomatic index (p > 0.05). We tested the association between bacterial diversity and host traits using GLM models. Bacterial diversity differed significantly between populations (LRchisq = 10.30, df=1, p=0.001; Figure 2), whereas no significant associations were detected with host traits length_centered, parasite load or sex. Due to the pronounced effect of host population, we tested each host trait individually by population. None of the host traits showed a significant correlation with Shannon diversity, as indicated by the Spearman rank correlation test (p > 0.05). We found no significant association between Fulton condition and overall bacterial diversity. However, in Bornholm, Simpson’s diversity index showed a significant positive correlation with hepatosomatic index, indicating that fish with relatively larger livers for their body size harbored higher bacterial diversity (rs = 0.37, df = 32, p = 0.036; see Figure S3). RDA analyses showed that sampling region has a strong effect on bacterial composition (F = 2.76, p = 0.005). Neither fish length (F = 0.77, p = 0.706), liver worm burden (F = 1.18, p = 0.249) nor sex (F = 0.95, p = 0.468) significantly explained variation in the microbial community structure (Figure S4). Given the strong regional effect, we subsequently assessed the independent contributions of fish length, liver worm count, and sex to microbial composition. First, we fitted a simplified RDA model that included only population as a fixed predictor, which explained 15.93% of the variation in community composition (F = 10.80, p = 0.001; Figure S4). In comparison, the full model attributed 15.95% of the variation to population (F = 2.76, p = 0.005). Variation partitioning analysis RDA revealed that population and host traits collectively accounted for 14.4% of the variation in bacterial community composition. The unique contribution of population was 2.7%, whereas the shared fraction (11.7%) represents the variation simultaneously explained by both population and host traits. Among the variables tested sex explained the largest proportion of variance (2.4%; RDA3 and RDA2) and showed a trend toward significance in Åland location (F = 1.59, p = 0.069, Figure S4), but not in Bornholm (F=0.66, p=0.73, Figure S3). When testing for interactions between populations and other host-related variables (fish length, number of liver worms and sex), redundancy analysis indicated that the interaction between sampling region and fish length had a marginally non-significant effect on bacterial composition (Df = 1, Variance = 0.01599, F = 1.63, p = 0.053). Interactions between populations and worm abundance or sex were not significant (p >0.05).

### Differentially abundant taxa

Using the LindA approach, we identified 889 ASVs that were differentially abundant between Bornholm and Åland, while DESeq2 detected 989 such ASVs. A total of 626 ASVs were identified by both methods, and these shared ASVs were used for subsequent downstream analyses. Among all detected phyla, only three—Actinobacteria (*Dermacoccaceae*), Firmicutes (f; *Planococcaceae*), and Proteobacteria (classes *Alphaproteobacteria, Betaproteobacteria*, and *Gammaproteobacteria*)— contained ASVs that were more abundant in Bornholm compared to Åland. Within the *Alphaproteobacteria* class, the genus *Rickettsia* was particularly more abundant in Bornholm, while *Polynucleobacter*, from the family *Burkholderiaceae* (*Betaproteobacteria*), also showed higher abundance in Bornholm. The *Gammaproteobacteria* group exhibited the most pronounced differences between Åland and Bornholm, with several genera displaying higher relative abundance in Bornholm (g; *Acitenobacter*, g; *Pseudomans* and g; *Pshycrobacter*, Figure 4; Figure S5). Several bacterial phyla were detected exclusively in Åland. Within the Proteobacteria phylum, ASVs affiliated with the classes Alphaproteobacteria (f. *Rhodobacteraceae*), Betaproteobacteria (*f. Comamonadaceae*), and Gammaproteobacteria (*f. Xanthomonadaceae* and f. *Moritellaceae*) were only present in Åland, among others. Within the Actinobacteria phylum, the class Thermoleophilia (*f. Thermoleophilaceae*) occurred exclusively in Åland. Most taxa within the class Actinobacteria (e.g., *f. Acidimicrobiaceae, f. Brevibacteriaceae, f. Dermabacteriaceae*) were also exclusively found in Åland. Within the Firmicutes phylum, the classes Clostridia (*f. Clostridiaceae, f. Ruminococcaceae, and f. Lachnospiraceae*) and Erysipelotrichia (*f. Erysipelotrichaceae*) were confined to Åland. In addition, within the class Bacilli, the families *Bacillales, Carnobacteriaceae*, and *Staphylococcaceae* were differentially abundant, presenting higher ASVs in Åland. (Figure 4).

**Figure 4.**
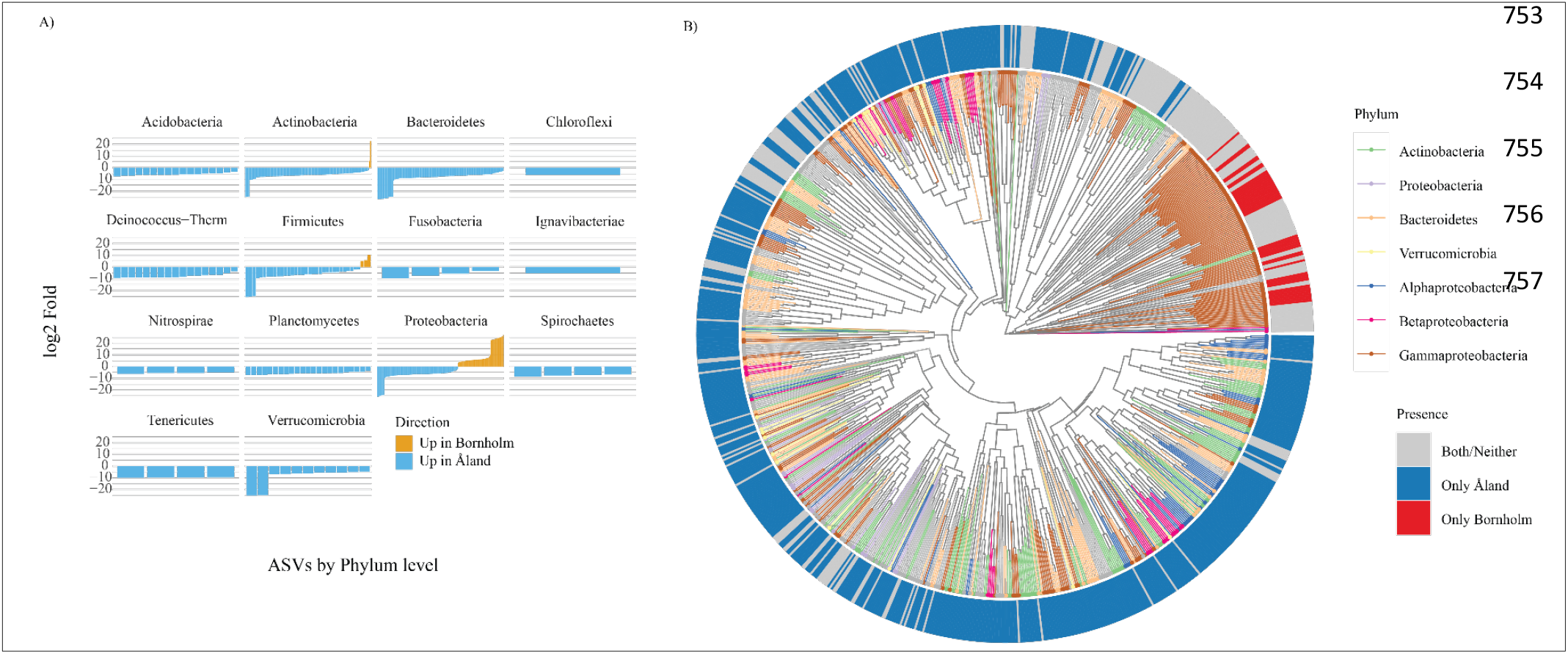
Differential abundance and phylogenetic distribution of gill microbiome ASVs in Eastern Baltic cod from Bornholm and Åland, identified using the conservative DESeq2 model. (A) Log2 fold change plots of ASV abundance by phylum, showing ASVs significantly higher in Bornholm (orange) or Åland (blue). (B) Phylogenetic tree of ASVs colored by phylum, with outer rings indicating ASV presence unique to Åland (blue), unique to Bornholm (red), or found in both/neither locations (grey).

### Prediction of differentially activated KO metabolic pathways

We used LindA and Deseq2 to estimate the differentially abundant KO metabolic pathways that are more activated between Åland and Bornholm. The most conservative approach was selected for visualization (Deseq2: N_(KO)_=2405). We grouped the KEGG Orthology (KO) metabolic pathways into 35 broad categories. While most metabolic pathway categories showed representation in both locations, the nervous system was exclusively associated with ASVs more abundant in Bornholm (Figure 5). In total, 50% of the metabolic pathways were differentially activated between Bornholm and Åland individuals, with lower activation observed in the Bornholm group: 28% were absent or significantly reduced, including immune disease, eukaryotic community and signaling pathways, while an additional 22% showed decreased activation compared to Åland. To ensure analytical robustness, both approaches were integrated, and only the KEGG Orthologs (KOs) identified by both methodologies were retained for downstream analysis. In total, 1,903 shared KOs were detected, representing the consensus set of metabolic pathways common to both approaches. The functional profiles of these KOs are presented in Figure 5S. Based on this subset, 24 metabolic pathways were predicted to be functionally active in Åland. In contrast, only two pathways—antimicrobial resistance and membrane transport—approached the threshold for functional activation in individuals from Bornholm (Figure S6).

**Figure 5.**
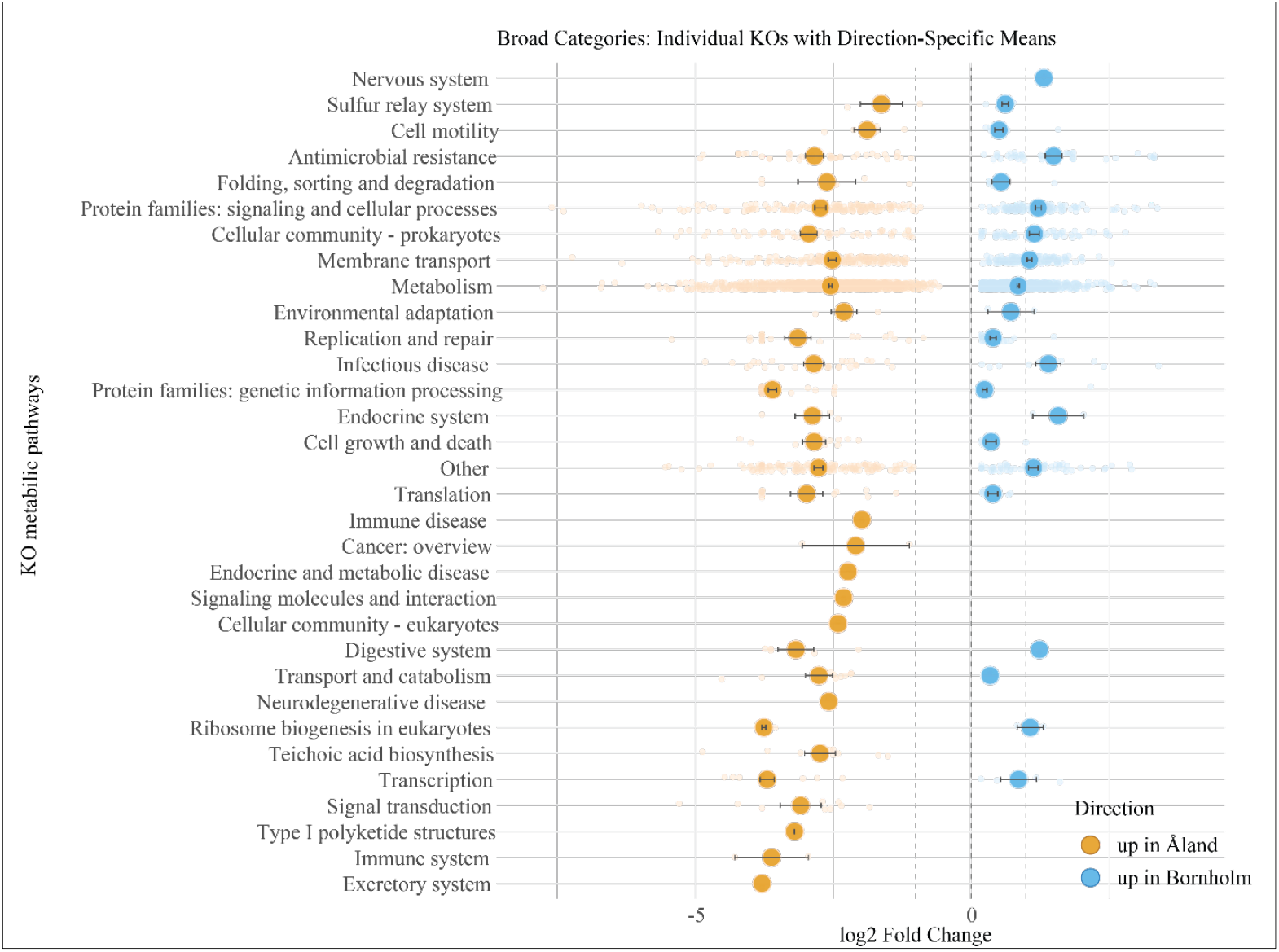
Differential abundance of KEGG Orthology (KO) metabolic pathways between Åland and Bornholm cod gill microbiomes. Each row represents a broad KO pathway category, with individual KOs plotted by log2 fold change. Orange circles indicate pathways upregulated in Åland, while blue circles indicate pathways upregulated in Bornholm. Circle size represents the magnitude of change, and horizontal bars denote the standard error for each category mean.

## Discussion

We characterized the diversity and composition of the gill-associated microbiota in two populations of eastern Baltic cod. We assessed differential bacterial abundance between the two regions and inferred GO metabolic pathways. Furthermore, we investigated whether gill microbiota composition is associated with specific host traits, including weight, length, liver weight, liver worm burden, and sex. This is the first study to characterize gill-associated microbiota in a Baltic fish species, providing novel insights into host–microbe–environment interactions in this ecosystem. Our results indicate considerable differences in bacterial diversity and community composition between cod from the two populations, with higher diversity observed in the Åland cod. This variation was driven primarily by population effects, and was not explained by host-specific traits. Contrary to expectations, no association was detected between liver worm burden and the gill-associated microbiota, despite the assumption that greater microbial diversity in uninfectied individuals may be indicative of improved host health. However, gill-associated microbiota was significantly linked to fish sex in Åland, with higher infection levels observed in smaller individuals from Bornholm as compared to Åland Sea. We also found that the phylum Proteobacteria exhibited the strongest difference in relative abundance between the two regions. Other phyla, such as Verrucomicrobia and Deinococcus–Thermus, were absent in Bornholm. Furthermore, we detected strong differences in GO metabolic pathways between the Åland and Bornholm regions. Specifically, KO pathways related to the nervous system were uniquely activated in Bornholm cod, potentially reflecting exposure to environmental stress or disease-related processes in this population (Toubanaki et al., 2022). A greater number of KEGG Ortholog (KO) pathways were active in Åland compared to Bornholm, indicating that individuals from Åland possess a wider functional repertoire of metabolic pathways. This enhanced metabolic capacity may confer increased resilience to environmental stressors, including coping with parasites (Sokolova, 2013, Liu et al., 2019).

Our results showed higher gill-associated alpha bacterial diversity in Åland as compared to cod from Bornholm Basin. This was contrary to our predictions as we expected a lower bacterial diversity towards higher latitude (Zhang et al., 2022, Fuhrman et al., 2008, Härer & Rennison, 2023). While some studies report higher bacterial diversity at lower latitudes (warmer temperatures near the equator), consistent with predictions of a latitudinal diversity gradient (Lear et al., 2017), others have observed no such relationship(Cai et al., 2024, Moss et al., 2020). Consistent with the latter studies, our data show a statistically significant yet reversed latitudinal trend in microbial diversity—increased diversity at higher latitudes—based on analyses of two populations. In addition, higher microbial diversity has been associated with increased host survival against viral pathogens in amphibians and fish, for example, 63% more bacterial taxa were detected in the gut of high-fitness as compared to low-fitness individuals in sticklebacks (Harrison et al., 2019). Therefore, the higher alpha diversity observed in our study may suggest that Baltic cod individuals from Åland are overall in better condition and exhibit greater fitness compared to those from the Bornholm Basin.

In addition to differences in alpha diversity, we observed distinct and statistically significant variation in bacterial community composition between the two populations. The composition of vertebrate skin microbiomes is typically shaped by multiple factors, including sex, diet, maternal effects, environmental conditions and geography (Ross et al., 2019). In salmon, skin-associated microbial communities were not correlated with the microbial composition of the surrounding water, despite experimental manipulation of salinity (Schmidt et al., 2015). In contrast, studies in several marine fish species have supported the idea that both seasonal variation and sampling location significantly influence microbial communities on fish skin (Liston, 1956, Ross et al., 2019, Larsen et al., 2015). Taken together, our findings suggest that variation between the populations in environmental variables, particularly salinity and dissolved oxygen, may play a central role in shaping microbiome structure in this system (Bierlich et al., 2018, Horsley, 1977). Host-specific skin compound characteristics and individual host traits may also play a significant role in modulating skin bacterial composition (Chiarello et al., 2015, Berggren et al., 2022). However, in accordance with previous studies (Chiarello et al., 2019, Uren Webster et al., 2018), we did not find a significant effect of host traits such as body size or weight on skin microbial community composition and diversity. Despite increasing interest in this field, fish skin microbiome remains comparatively understudied relative to the gut microbiome. Further research across Baltic fish species and populations is needed to clarify the potential contributions of geography, environmental factors, and host genetic background to variations in skin microbiome composition.

Several studies have investigated the factors influencing the health status of Eastern Baltic cod populations (Eero et al., 2023, Eero et al., 2020, Froese, 2025). One potential contributor are fishing practices that selectively remove larger individuals, thereby favoring smaller fish via selection processes (Han et al., 2025). Another important factor is infection by parasites, in the present case the parasitic liver worm. Eastern Baltic cod has been infected with *C. osculatum* infections over the years, and bioenergetic modelling and a controlled laboratory experiment suggests that growth rates are reduced in heavily infected individuals compared to non-infected ones (Ryberg, 2020, Ryberg et al., 2023). Dysbiosis of the fish gut microbiome has been reported in chub infected with helminths (Colin et al., 2022) and in common carp infected with intestinal tapeworms (Fu et al., 2019). Given the documented impact of parasitic infections on host-associated microbiota, we hypothesized that liver worm infections could similarly influence the gill-associated microbial community. However, we found no significant association between gill-associated microbiome composition and liver worm infection in the present study. Future research should prioritize metatranscriptomic analyses to detect functional microbiome shifts associated with infection that are not apparent from taxonomic profiles alone, and should integrate host-level holobiont approaches, such as host immune gene expression, to evaluate the direct effects of infection on microbial communities.

Our results show that Proteobacteria dominated the gill-associated microbiota in both cod from the Åland Sea and the Bornholm Basin, with Gammaproteobacteria phyla being the most differentially abundant class. This finding is consistent with observations from other marine fish species (Larsen et al., 2015, Wang WenWen et al., 2010, Bell et al., 2024), where Proteobacteria are consistently reported as the predominant phylum in fish skin and gut microbiomes (Berggren et al., 2022, Ross et al., 2019, Gomez & Primm, 2021, Krotman et al., 2020). Within the Gammaproteobacteria class, we found that Pseudomonas was significantly more abundant, consistent with earlier studies in marine fish in Bornholm (Larsen et al., 2015, Liu et al., 2024). Several strains of Pseudomonas are widely recognized as opportunistic fish pathogens, while others exhibit bioremediation and biocontrol activities (Aydin et al., 2023, Debnath et al., 2023). We also found that the genus Psychrobacter was abundant in Borhnolm site. Psychrobacter includes strains that act as opportunistic pathogens in both fish and humans, causing severe disease in marine fish in the Red Sea (El-Sayed et al., 2023). However, some Psychrobacter strains have also been reported as opportunistic symbionts with protective functions (Wuertz et al., 2023). Although this study is the first study on cod to identify a specific group of bacteria that warrant attention due to their potential serious impact on host health, definitive assignment of specific pathogenicity could not be made within this study.

Notably, the nervous system KEGG Orthology (KO) pathway exhibited significant activation exclusively in the Bornholm population using the most conservative analytical approach (DESeq2), which identified fewer differentially abundant Kos (N=2405) than the LindA method (N=5062). .. This activation may be related to hypoxic conditions prevalent in the Bornholm area (Fritsche & Nilsson, 1990), as previous studies have demonstrated that autonomic nervous system regulation of blood pressure and heart rate occurs during hypoxia in cod species, potentially triggering this pathway (Fritsche & Nilsson, 1990). Another potential cause of nervous system metabolic pathway activation could be prolonged exposure to toxins or anthropogenic pollutants, such as polycyclic aromatic hydrocarbons (PAHs) and perfluoroalkyl substances (PFAS), which are increasing in the Baltic Sea (Soerensen et al., 2024). These pollutants have been reported to affect neuro-transcriptomics in female Atlantic cod (Denzil Rozario, 2022). Additionally, nervous system activation may result from infectious diseases, such as viral nervous necrosis (VNN), which can alter the microbiome upon infection and cause focal vacuolating encephalopathy and retinopathy (Krasnov et al., 2013). The precise cause of nervous system pathway activation remains undetermined, as our study does not provide direct evidence; however, these findings serve as a foundation for further investigation.

In conclusion, significant differences in bacterial diversity and composition, and enrichment of disease-associated bacteria were observed in the Bornholm area compared to Åland, suggesting that the southern population of Eastern Baltic cod is in poorer health condition compared to the northern population in the Åland Sea. While many studies report pathogenicity, it is often strain-specific; therefore, microbiome surveys and KEGG Orthology (KO) metabolic pathway analyses based on 16S metabarcoding cannot directly determine pathogenic potential and have to be interpreted with caution (Wiles & Guillemin, 2019). To address the direct relationship between pathogenicity and body size in the Baltic cod, we highlight the urgent need for metagenomic approaches to more accurately assess strain-specific pathogenicity and to precisely infer metabolic pathway activation.

## Ethics approval and consent to participate

Not applicable

## Consent for publication

Not applicable

## Availability of data and materials

All data generated or analyzed during this study are included in this published article and its supplementary information files. The R scripts and raw data will be deposited in a public repository and made available upon publication.

## Authors’ contributions

**CCM** conceptualized the study, performed all wet-lab experiments, conducted data analyses, secured wet-lab funding, and wrote the initial manuscript draft. **AL** contributed to conceptualization, obtained primary funding for sampling, and revised the final manuscript draft. **HY** and **GOM** conducted field sampling and obtained some funding for the sampling. All authors reviewed and approved the final manuscript version.

## Acknowledgements

We gratefully acknowledge all participants involved in the SVEA (BITS) sample collection at Bornholm, as well as the private fishermen who contributed sampling from the Åland site.

