## SUPPLEMENTARY INFORMATION: FIGURES AND TABLES for "Gill-associated microbiome as indicators of population stress and condition in Eastern Baltic Cod"

### Supplementary figures

**Figure S1**. Taxonomic diversity of gill microbiomes in Eastern Baltic cod. (A) Overall distribution of bacterial genera by phylum, showing that Proteobacteria, Bacteroidetes, Actinobacteria, and Firmicutes dominate the genus-level diversity. (B) Taxonomic diversity by phylum and region, comparing the composition between Åland (orange) and Bornholm (blue). Percentages indicate the proportion of total genera within each phylum and region.

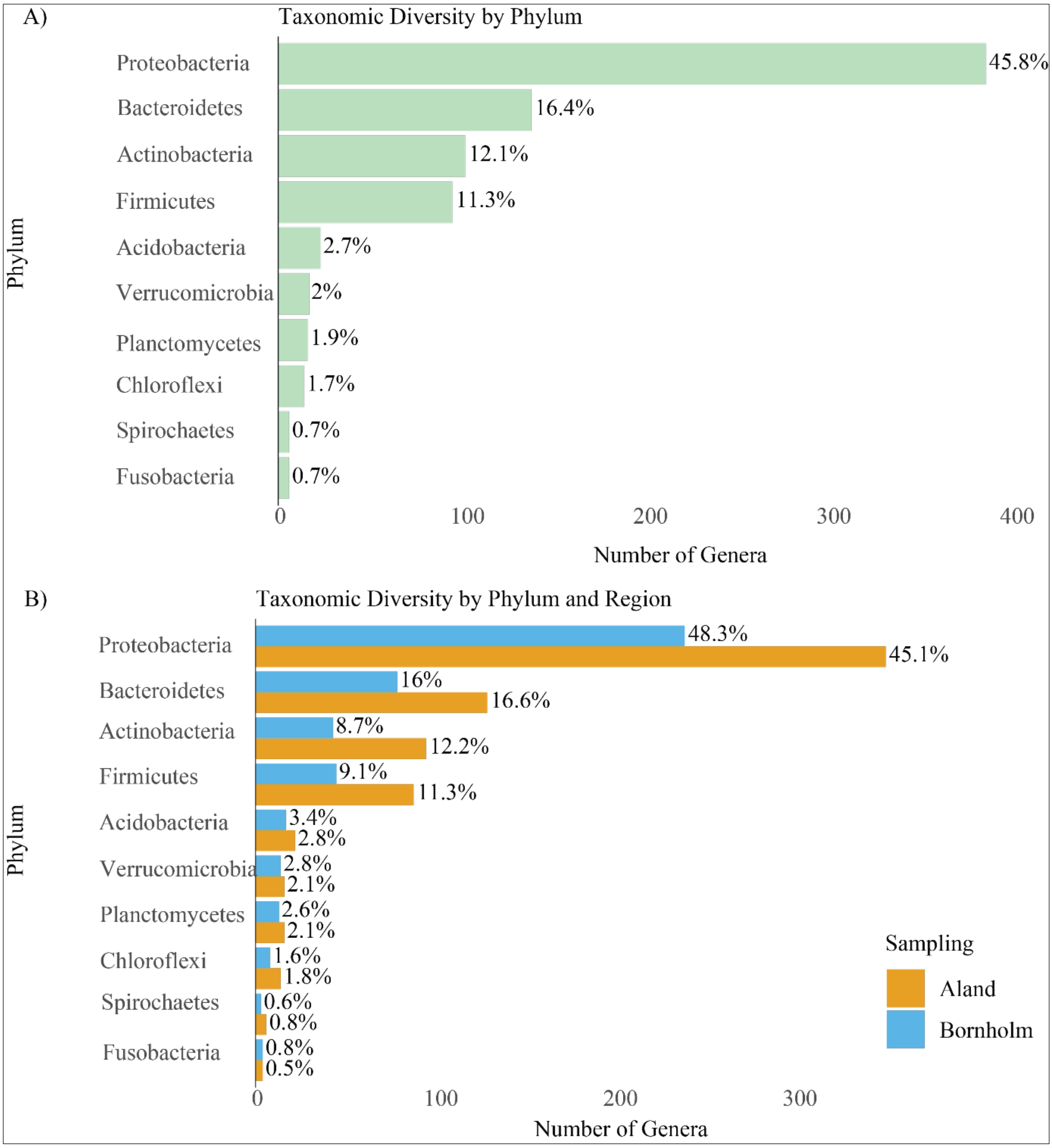

**Figure S2**. Liver worm burden in Eastern Baltic cod from Åland and Bornholm. (A) Boxplot of log-transformed liver worm counts ($p < 0.001$). (B) RDA ordination plots illustrate variation in worm burden patterns within each sampling site, with individual points colored according to log-transformed worm count. The color gradient reflects increasing worm burden from blue (low) to red (high).

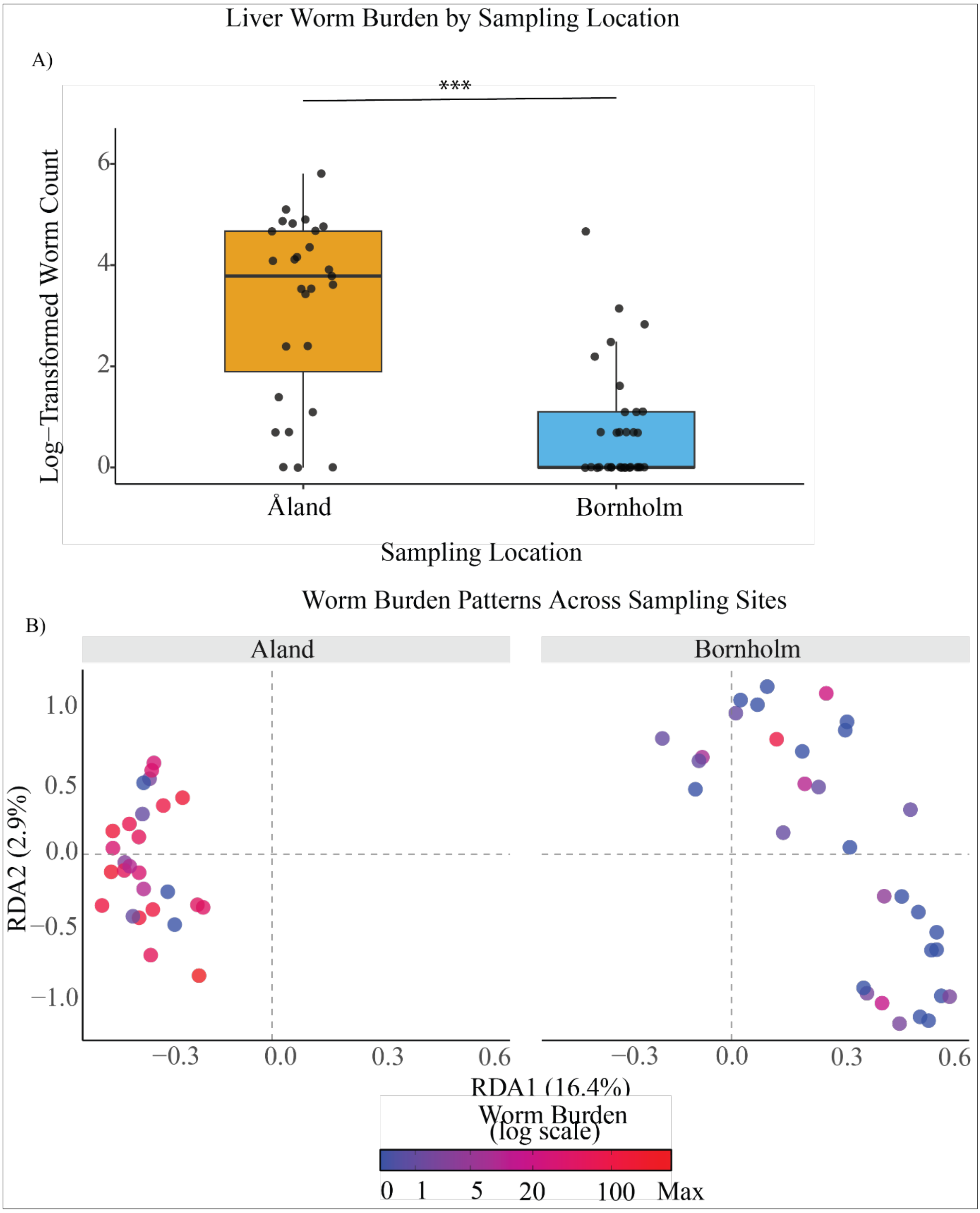

**Figure S3.** Spearman rank correlations between gill microbiome alpha diversity and hepatosomatic index in Baltic cod.

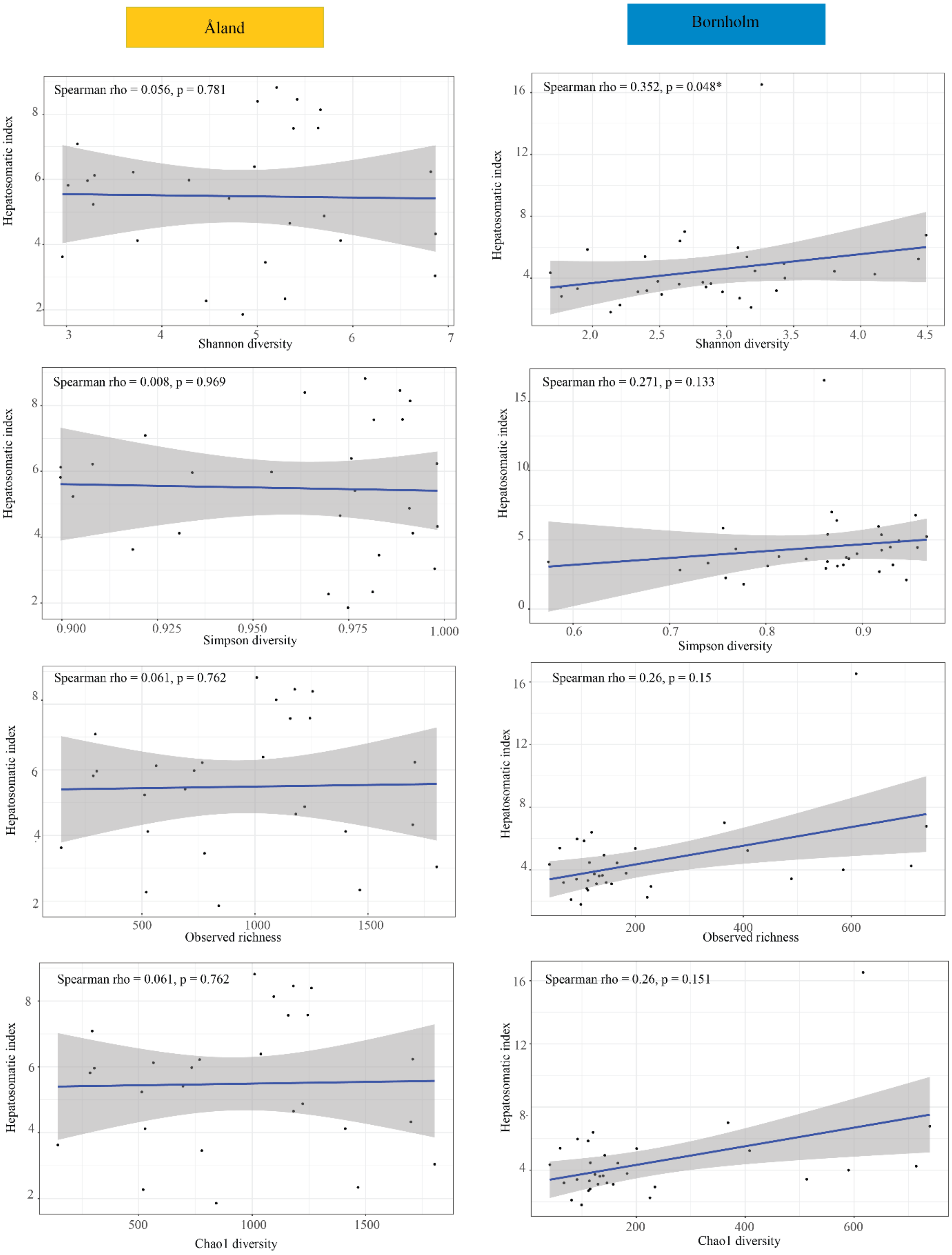

**Figure S4**. Redundancy analysis (RDA) of microbial community composition in Eastern Baltic cod gill microbiomes. (A) RDA plot showing the influence of fish length, sex, and liver worm count on microbial communities, with samples colored by sampling site (Bornholm in blue, Åland in orange). (B) Sex-based differences in microbiome composition at the Åland site, with samples colored by sex. (C) Sex-based differences in microbiome composition at the Bornholm site, with samples colored by sex. Percentage values on axes indicate the variation explained by each RDA component.

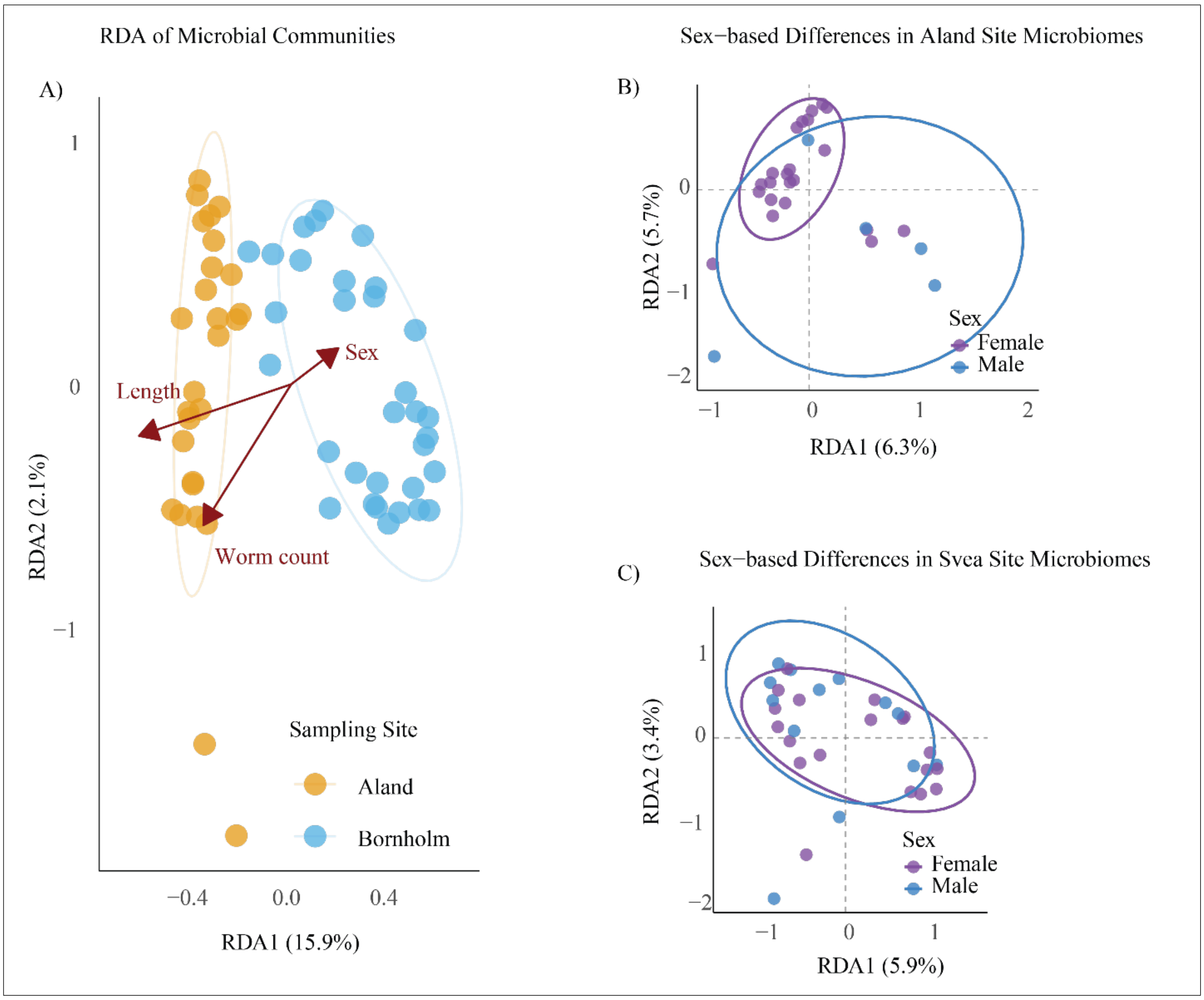

**Figure S5**. Distribution of ASVs within major families and genera of Gammaproteobacteria in Eastern Baltic cod gill-associated microbiomes by using the common ASVs generated by both methods LindA and Deseq2. The heatmap shows the number of ASVs (n_ASVs, color scale) detected for each genus–family combination in Bornholm.

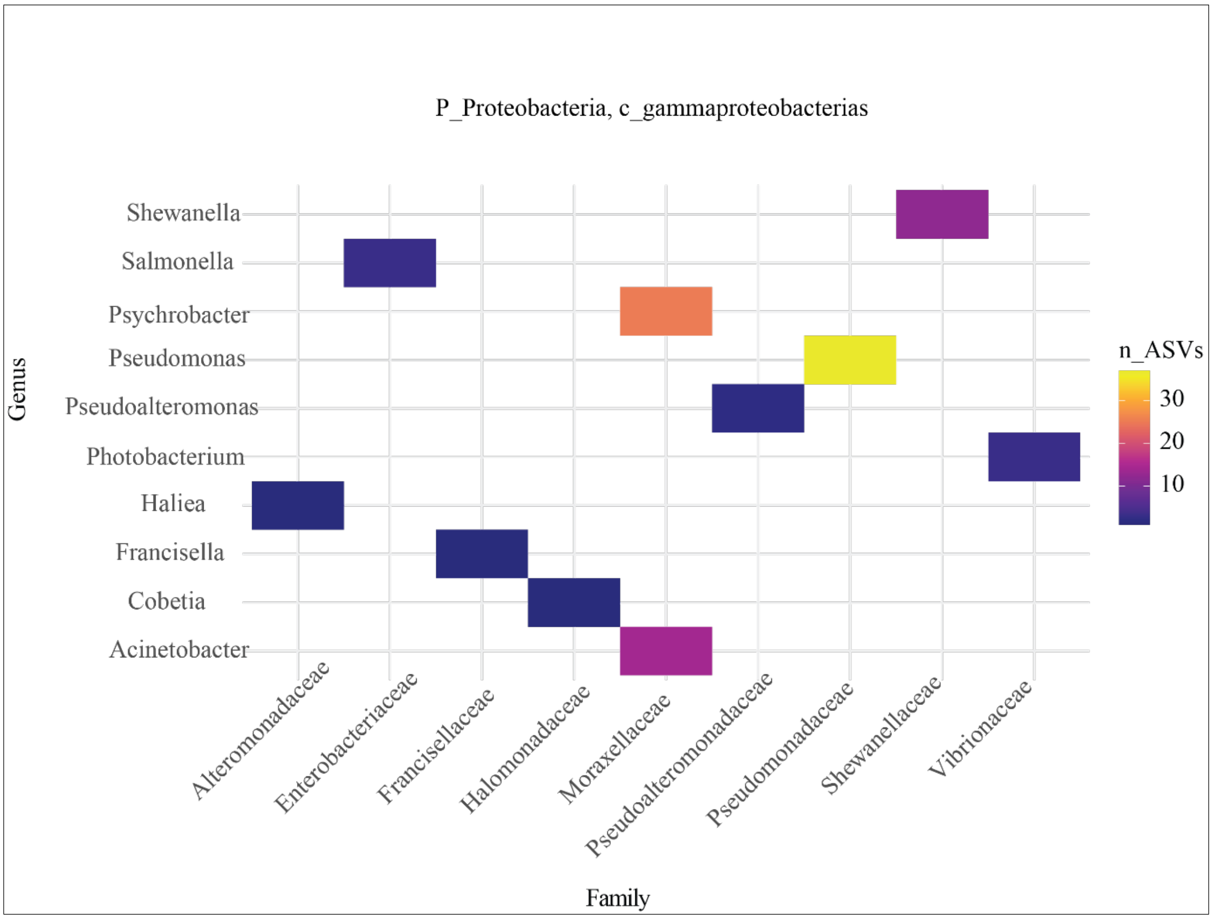

**Figure S6**. Comparison of KO metabolic pathway detection and differential abundance between LindA and DESeq2 models.(A) Venn diagram illustrating the overlap and unique KO metabolic pathways identified by LindA (blue) and the more conservative DESeq2 model (red), with 1,903 pathways shared, 3,159 detected solely by LindA, and 1,312 by DESeq2 alone. (B) Differential abundance analysis of broad KO pathway categories, showing log2 fold changes for individual KOs. Orange circles indicate pathways enriched in Åland, while blue indicates those enriched in Bornholm. Circle size reflects the magnitude of change, and horizonal lines denote standard error for each category mean.

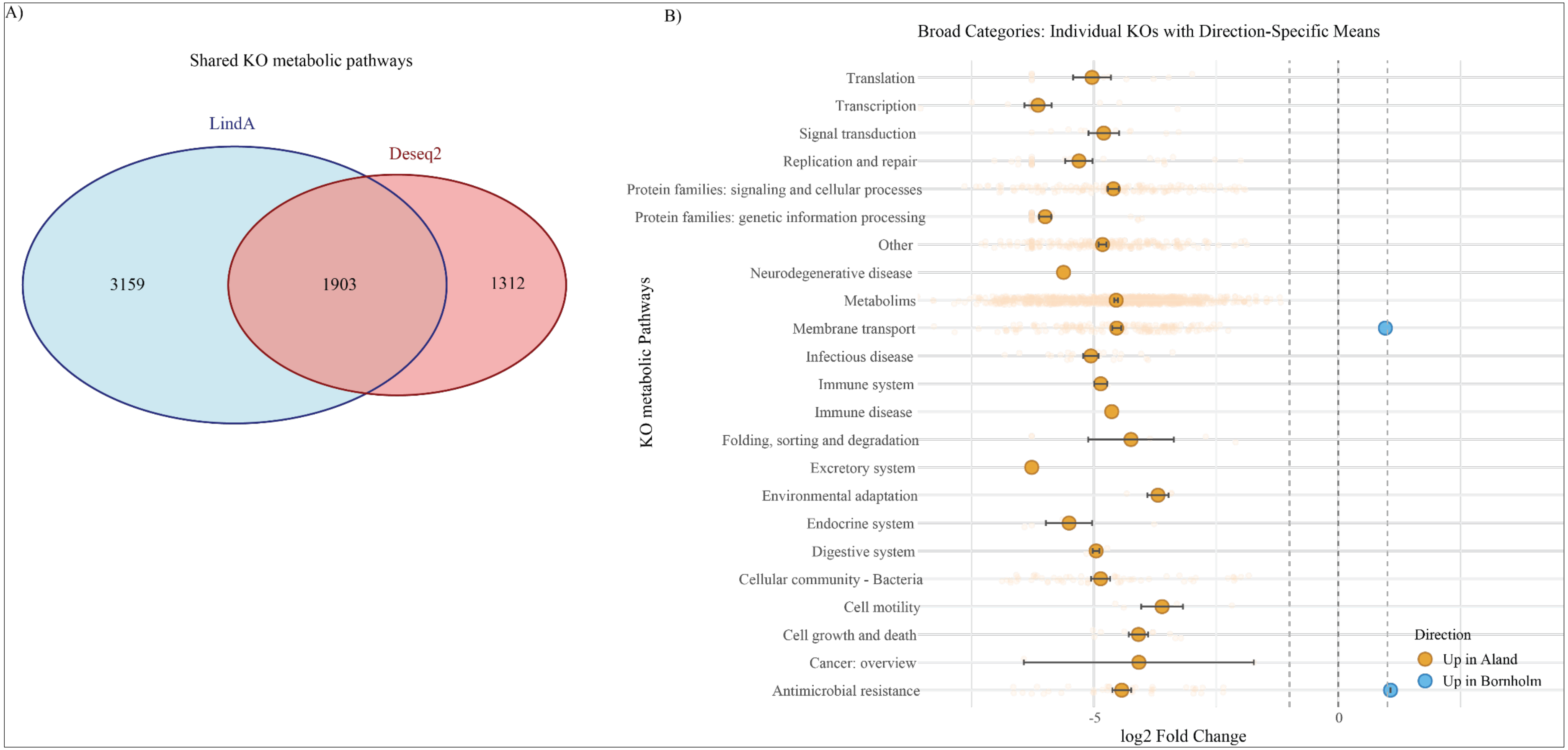
